# Molecular basis of product recognition during PIP5K-mediated production of PI(4,5)P_2_ with positive feedback

**DOI:** 10.1101/2023.09.04.556152

**Authors:** Benjamin R. Duewell, Katie A. Faris, Scott D. Hansen

## Abstract

The ability for cells to localize and activate peripheral membrane binding proteins is critical for signal transduction. Ubiquitously important in these signaling processes in eukaryotic cells are phosphatidylinositol phosphate (PIP) lipids, which are dynamically phosphorylated by PIP lipid kinases on intracellular membranes. Functioning primarily at the plasma membrane, phosphatidylinositol-4-phosphate 5-kinases (PIP5K) catalyze the phosphorylation of PI(4)P to generate most PI(4,5)P_2_ lipids found in cells. Recently, we determined that PIP5K displays a positive feedback loop based on membrane-mediated dimerization and cooperative binding to its product, PI(4,5)P_2_. Here, we examine how two PIP5K motifs contribute to PI(4,5)P_2_ recognition to control membrane association and catalysis. Using a combination of single molecule TIRF microscopy and kinetic analysis of PI(4)P lipid phosphorylation, we map the sequence of steps that allow PIP5K to cooperatively engage PI(4,5)P_2_. We find that the specificity loop regulates the rate of PIP5K membrane association and helps orient the kinase to more effectively bind PIP lipids. Attaching the PIP5K specificity loop to other peripheral membrane binding proteins can enhance their membrane binding dynamics. After correctly orienting on the membrane, PIP5K transitions to binding PIP lipids in a structural motif previously referred to as the substrate or PIP binding motif (PIPBM). Our data reveals that the PIPBM has broad specificity for anionic lipids and serves a critical role in regulating membrane association in vitro and in vivo. The strength of the interaction between the PIPBM and PIP lipids depends on the membrane density and the extent phosphorylation on the inositol head group. We propose a two-step membrane binding model where the specificity loop and PIPBM act in concert to help PIP5K orient and productively engage anionic lipids to drive the positive feedback during PI(4,5)P_2_ production.

## INTRODUCTION

Phosphatidylinositol phosphate (PIP) lipids are important second messengers that function as molecular scaffolds for numerous membrane binding proteins. PIP lipids are dynamically interconverted between seven different species by specialized kinases and phosphatases (Balla 2013). Research has demonstrated that each PIP lipid species exists in spatially regulated subcellular membranes, functioning to recruit specific enzymes to their necessary location for proper signaling (Hammond and Burke 2020). The most abundant PIP lipid that mediates signaling at plasma membrane is phosphatidylinositol-4,5-bisphosphate (PI(4,5)P_2_), comprising between 2-5% of total plasma membrane lipids (Mitchell, Ferrell, and Huestis 1986; Nasuhoglu et al. 2002; Wenk et al. 2003). Unlike most other PIP lipids, PI(4,5)P_2_ is omnipresent at the plasma membrane and is critical for continual execution of many basic cell functions (Di Paolo and De Camilli 2006). These signaling events include cytoskeletal assembly, ion channel gating (Hansen 2015; Huang, Feng, and Hilgemann 1998), and endocytosis (Antonescu et al. 2011). Without constantly available pools of PI(4,5)P_2_, cellular signaling events are hindered and cells adopt disease-like states (Mandal 2020). The requirement of the cell to have relatively constant PI(4,5)P_2_ levels suggests that the enzyme(s) capable of synthesizing this lipid should be capable of activity with little regulation.

Phosphatidylinositol-4-phosphate 5-kinase (PIP5K) is the enzyme responsible for synthesizing the bulk of PI(4,5)P_2_ lipids in eukaryotic cells (Rameh et al. 1997; Wills and Hammond 2022). PIP5K enzymes function by binding their substrate, phosphatidylinositol-4-phosphate (PI(4)P), then catalyzing the phosphorylation of the 5-hydroxyl on the inositol lipid head group in the presence of Mg-ATP. This activity has been described as specific to the PI(4)P substrate, although PIP5K can also phosphorylate the 5-OH of other PIP lipid species with lower efficiency (Kunz et al. 2000, 2002; Tolias et al. 1998; Zhang et al. 1997). In mammals, PIP5K exists as three paralogs (α, β, and γ) that display tissue specific expression patterns (Ishihara et al. 1996, 1998; Loijens and Anderson 1996). Localization studies in macrophages have shown that PIP5K paralogs predominately localize to the plasma membrane (Fairn et al. 2009). PIP5K has been implicated in many cellular pathways, with specific regulatory importance in the Wnt pathway (Pan et al. 2008), phagocytosis (Coppolino et al. 2002; Mao et al. 2009), and focal adhesions (Di Paolo et al. 2002; Ling et al. 2002). More recently, medical studies have begun to target PIP5K with high potential as a cancer therapeutic (Burke et al. 2023). For example, studies that have knocked out PIP5K from both prostate and breast cancers models showed diminished tumorigenesis and metastatic properties. The drugs designed to modulate PIP5Kα activity in these models has incredibly promising effects on tumor growth and proliferation, but current inhibitors have low specificity for PIP5K. As our understanding of PIP5K’s role in tumor proliferation improves, it is important that we continue to decipher the molecular mechanisms that regulate PIP5K membrane localization and activity in cells.

Recently, our lab has employed supported lipid bilayers (SLBs) and Total Internal Reflection Fluorescence Microscopy (TIRF-M) to directly visualize PIP5K membrane binding dynamics with single molecule resolution. We discovered that human PIP5KB binds cooperatively to its product, PI(4,5)P_2_ (Hansen et al. 2019), which enhances PIP5K membrane localization through a PI(4,5)P_2_-mediated positive feedback loop. When PIP5K crosses a threshold membrane surface density it can also dimerize, which potentiates lipid kinase activity (Hansen et al. 2022). This suggests that the site of PI(4,5)P_2_ binding may be an important step for plasma membrane localization in cells, as PI(4,5)P_2_ lipids exist almost exclusively at the plasma membrane. Currently, the site of high affinity PI(4,5)P_2_ binding is unknown.

To date, much work has been done to define PIP5Ks enzymatic activity, including identifying key motifs important for substrate specificity, ATP binding, and hydrolysis (Kunz et al. 2002; Liu et al. 2016; Muftuoglu et al. 2016). Structural biochemistry has helped elucidate the mechanism controlling PIP5K substrate recognition (Liu et al. 2016; Muftuoglu et al. 2016). X-ray crystallography of zebrafish PIP5K (zPIP5K) determined that PIP5K binds to its substrate, PI(4)P, through the DLKGSxxxR motif, which is also referred to as the PI(4)P or substrate binding motif (Hu et al. 2015; Muftuoglu et al. 2016). Mutations in the PI(4)P binding motif of PIP5Ks abolishes lipid kinase activity (Muftuoglu et al. 2016). PIP5K substrate specificity has been found to be mediated by a structural motif referred to in the literature as either the activation loop or referred to here as the specificity loop. Structural analysis of zPIP5Kα using nuclear magnetic resonance (NMR) suggests that the specificity loop exchanges between a disordered and an alpha helical conformation (Liu et al. 2016). Membrane-lipid interactions are hypothesized to shift the equilibrium to favor the alpha helical conformation of the specificity loop, allowing hydrophobic residues in the amphipathic helix to insert into a lipid bilayer (Liu et al. 2016). How the specificity loop regulates PIP lipid interactions, however, remains unknown. While these studies have revealed which motifs are critical for PIP5K lipid kinase activity, we do not currently understand how these various properties contribute to the cooperative PI(4,5)P_2_ binding, membrane mediated dimerization, and positive feedback. Deciphering these mechanisms could help explain why PIP5K is predominantly localized and activated at the plasma membrane.

Here we present a mutational analysis and single molecule TIRF-M study of PIP5K to determine the sequence of molecular interactions that control PIP5K membrane docking and PI(4,5)P_2_ engagement. The initial membrane interaction is regulated by the specificity loop, which plays an important first step for membrane binding but is not essential for productive membrane docking. We determined that the mechanism of substrate specificity using a series chimera proteins containing specificity loop to show that this initial binding step of PIP5K is specific to PI(4) P lipids, defining the molecular mechanism of substrate specificity previously unknown. Interactions with the lipid product, PI(4,5)P_2_, were paradoxically mediated through region of the kinase referred to as the PI(4) P binding motif (PIPBM) or substrate binding motif. We further identified that the PIPBM is a broad anionic lipid sensor that can associate with lipids including phosphatidylserine, PI(4,5)P_2_, and PI(3,4,5)P_2_. Overall, the strength of lipid interactions with the PIPBM were strongly correlated with the valency of phosphorylation on the PIP lipid head group, with triply phosphorylated PIP lipids binding strongest. Supporting that PIPBM controls membrane localization in addition to catalysis, we find that PIPBM mutants can still phosphorylate PI(4)P when artificially membrane localization. Finally, we used new insight about PIP5Ks membrane binding mechanism to isolate separation of function mutants that are catalytically dead, but retain their ability to membrane localize like the wild type kinase. Overall, this work improves our understanding of the molecular mechanisms underlying PIP5K membrane targeting and specificity.

## RESULTS

### The specificity loop PIP5K localization to PI(4,5)P_2_ containing membranes

Previous studies indicate that PIP5K substrate recognition is regulated by the specificity loop (Kunz et al. 2000, 2002; Liu et al. 2016; Muftuoglu et al. 2016). Positioned within ∼20Å of the active site, the specificity loop is thought to help organize the active site to orient substrate, PI(4)P, for catalysis (Muftuoglu et al. 2016). Nuclear magnetic resonance (NMR) experiments also suggest that membrane docking of PIP5K causes the unstructured specificity loop to adopt an alpha helical conformation, which reportedly inserts into lipid membranes (Liu et al. 2016). Membrane insertion of an amphipathic helix is predicted to significantly enhance the membrane dwell time of PIP5K and could contribute to the positive feedback mechanism observed during PI(4,5)P_2_ production (Hansen et al. 2019). To determine how the specificity loop contributes to PIP5K localization and positive feedback, we disrupted alpha helix formation in the specificity loop by introducing a W365P mutation in PIP5KB (**Figure 1A**). A homologous mutation (i.e. W393P) was previously shown to reduce catalytic activity of zebrafish PIP5KA 25-fold relative to the wild type kinase (Liu et al. 2016). Consistent with previous observations, PIP5KB (W365P) displayed a 50-fold reduction in lipid kinase activity compared to wild type PIP5KB when measured on supported lipid bilayers (SLBs) using TIRF microscopy (**Figure 1B**). To determine the extent to which the reduced activity of PIP5KB (W365P) is due to defects in membrane localization, we used sortase mediated peptide ligation to fluorescently label PIP5KB with Alexa Fluor 647 (AF647) and visualized the mutant kinase membrane localization. Unlike wild type and dimerization deficient (i.e. D51R) AF647-PIP5KB, localization of AF647-PIP5KB (W365P) did not exhibit cooperative membrane recruitment that was coupled to production of PI(4,5)P_2_ lipids (**Figure 1C**). Overall, the reduced PIP5KB (W365P) activity was strongly correlated with a dramatic decrease in membrane localization.

**Figure 1.**
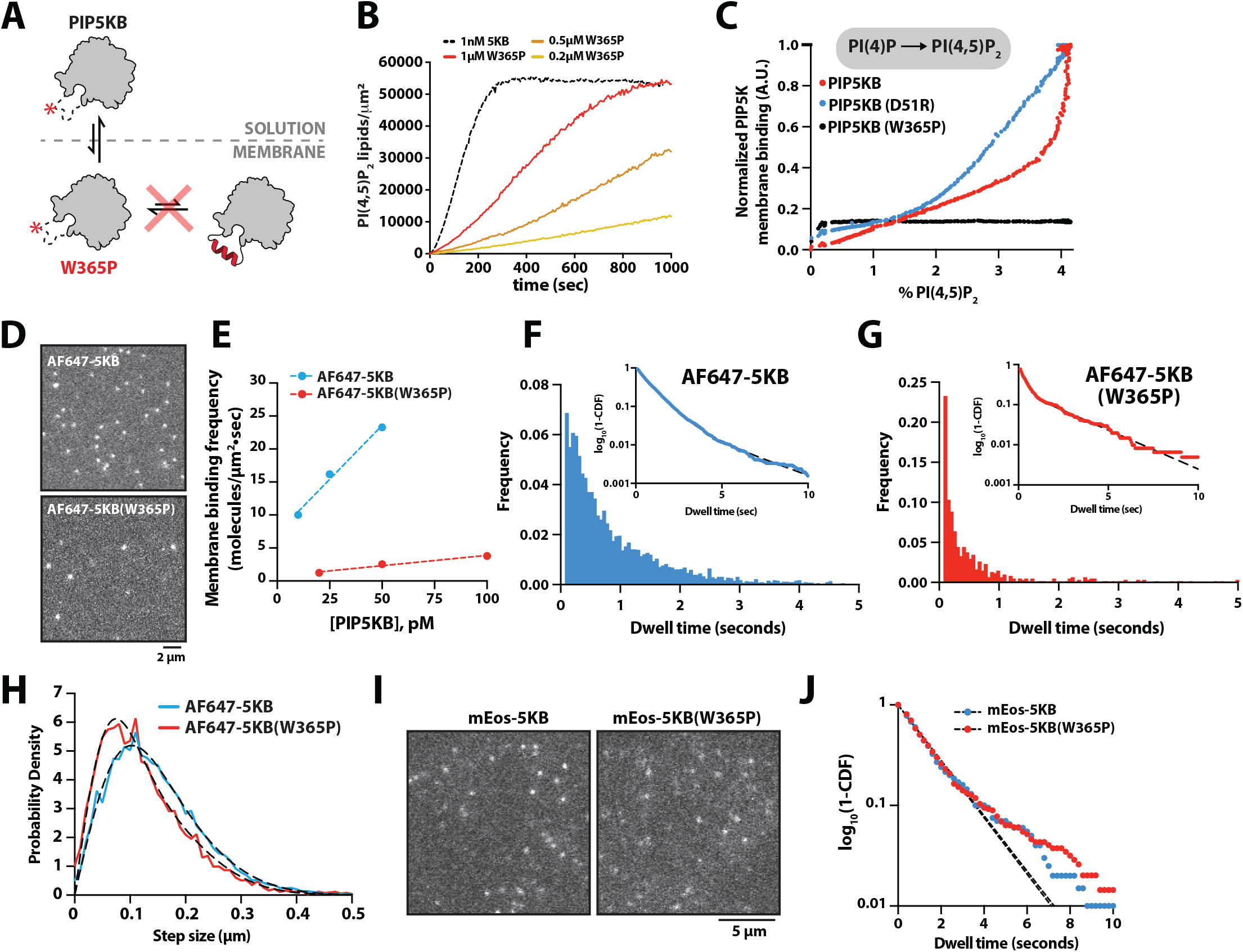
The specificity loop regulates the membrane association and dissociation rate constants. **(A)** Cartoon illustrating how the W365P mutation disrupts the specificity loop alpha helical conformation in PIP5KB. **(B)** Lipid kinase activity measured in the presence of PIP5KB and PIP5KB (W365P). Production of PI(4,5)P_2_ was monitored in the presence of 20 nM Cy3-PLCδ and the indicated concentration of PIP5KB. **(C)** PIP5KB (WT and D51R) but not PIP5KB (W365P) display cooperative PI(4,5) P_2_ dependent changes in supported membrane localization. Phosphorylation of PI(4)P and membrane localization of AF647-PIP5KB were monitored in the presence of 5 nM AF647-PIP5KB (WT or mutant) and 20 nM Cy3-PLCδ. **(B-C)** Initial membrane composition: 96% DOPC, 4% PI(4)P. **(D)** Representative TIRF-M images showing membrane localization in the presence of 100 pM AF647-PIP5KB or AF647-PIP5KB (W365P). **(E)** Experimental measure of membrane binding frequency (*k*_*ON*_) in the presence of 10-50 pM AF647-PIP5KB and 20-100 pM AF647-PIP5KB (W365P). **(F-G)** Single molecule dwell time distributions measured in the presence of either AF647-5KB or AF647-5KB(W365P). **(H)** Step size distribution comparing the membrane diffusivity of AF647-5KB and AF647-5KB(W365P). **(D-H)** Membrane composition: 96% DOPC, 4% PI(4,5)P_2_. **(I)** Representative TIRF-M images of mEos3.2-PIP5KB and mEos3.2-PIP5KB(W365P) plasma membrane localization following photoconversion with 405 nm light in HEK293T cells. **(J)** Single molecule dwell time distributions of mEos3.2-PIP5KB and mEos3.2-PIP5KB (W365P).

Since membrane binding of AF647-PIP5KB (W365P) was unresponsive to the generation of PI(4,5)P_2_ lipids, we hypothesized that the specificity loop could directly sense PI(4,5)P_2_ lipids, in addition to playing a critical role in substrate recognition. When we compared the single molecule localization of AF647-PIP5KB and AF647-PIP5KB (W365P) we observed a striking difference in the total number of membrane binding events on PI(4,5)P_2_ containing bilayers (**Figure 1D**). To determine the molecular basis of this difference, we measured the single molecule membrane binding frequency of AF647-PIP5KB and AF647-PIP5KB (W365P). Mutating the specificity loop resulted in a 10-fold decrease in the association rate constant (*k*_*ON*_) compared to wild type PIP5KB (**Figure 1E**). In addition, to the observed reduction in *k*_*ON*_ for AF647-PIP5KB (W365P), disrupting the specificity loop slightly reduced the single molecule dwell time in the presence of PI(4,5)P_2_ (**Figure 1F-1G**). This was surprising considering that the specificity loop has been proposed to penetrate directly into membranes (Liu et al. 2016), which is predicted significantly impact the dwell time of peripheral membrane binding proteins. Quantification of membrane diffusivity revealed that disruption of the specificity loop, also subtly changed in the diffusion of AF647-PIP5KB (W365P) compared to wild type AF647-PIP5KB (**Figure 1H**). Together, these findings suggest that while binding frequency and dwell time were diminished in the absence of the specificity loop, but PIP5K (W365P) can still strongly associate with PI(4,5)P_2_ lipids.

To determine whether membrane binding of PIP5KB (W365P) was similarly altered in vivo, we established conditions to visualize the plasma membrane localization of PIP5KB in HEK293T cells with single molecule resolution using TIRF-M. For these experiments, genes encoding either mEos3.2-PIP5KB or mEos3.2-PIP5KB (W365P) were transiently expressed in HEK293T cells under a heat shock promoter, which yielded a near single molecule membrane density of fluorescently tagged kinase localized to the plasma membrane (**Figure 1I**). Like our in vitro measurements on SLBs, mEos3.2-PIP5KB and mEos3.2-PIP5KB (W365P) displayed nearly identical dwell times in vivo (**Figure 1J**). The frequency of observing membrane localization of mEos3.2-PIP5KB (W365P), however, was greatly diminished. Together, these data indicate that the specificity loop facilitated PI(4,5)P_2_ binding but is not essential. Most important, the specificity loop serves a critical role in PIP5K substrate or product recognition during the initial membrane docking step.

### Chimeric proteins containing the PIP5K specificity loop display enhanced PIP specificity

Previous research indicates that the specificity loop controls membrane docking of PIP5K through substrate specificity (Kunz et al. 2000; Liu et al. 2016; Muftuoglu et al. 2016). This is supported by molecular dynamic simulations that indicate membrane docking relies on the initial engagement of PI(4)P by the specificity loop, which then transitions to the active site for catalysis (Amos et al. 2019). Based on our single molecule dwell time analysis of AF647-PIP5KB on SLBs, the specificity loop regulates the rate of membrane association (*k*_*ON*_) on PI(4,5)P_2_ containing membranes. Given the proximity between the specificity loop and active site of PIP5K, it’s thought that PIP lipids can associate with only one of these structural features at a time. To determine whether the specificity loop can function as a general PI(4)P and/ or PI(4,5)P_2_ lipid sensor in the absence of the entire PIP5K protein, we created a chimeric mNeonGreen (mNG) fusion protein containing the PIP5K specificity loop fused to the pleckstrin homology (PH) domain derived from phospholipase C-δ1 (i.e. mNG-5K(SL)-PLCδ) (**Figure 2A**). Characterization of the mNG-5K(SL)-PLCδ membrane binding properties in the context of PLCδ provided an approach for measuring the potential gain in PIP lipid specificity and enhanced membrane binding affinity due to the addition of the PIP5KB specificity loop. Using single molecule TIRF-M we examined whether mNG-5K(SL)-PLCδ acquired membrane binding specificity for either PI(4) P or PI(5)P lipids (**Figure 2B**). Consistent with PH-PLCδ lacking specificity for either PI(4)P or PI(5)P, mNG-PLCδ did not strongly interact with either lipid on supported membranes (**Figure 2B**). By contrast, we observed an increase in the single molecule dwell time for mNG-5K(SL)-PLCδ in the presence of PI(4)P, but not PI(5)P (**Figure 2B**). This result supports that the PIP5K specificity loop can discriminate between PI(4) P and PI(5)P, even when fused to a different peripheral membrane binding protein. Note that we also attempted to visualize membrane interactions of mNG fused to the PIP5K specificity loop (i.e. mNG-5K-SL), however, this chimeric protein displayed very transient membrane interactions that were not quantifiable. Presumably, the additional weak electrostatic interactions provided by PH-PLCδ facilitate membrane association in the presence of 5K(SL).

**Figure 2.**
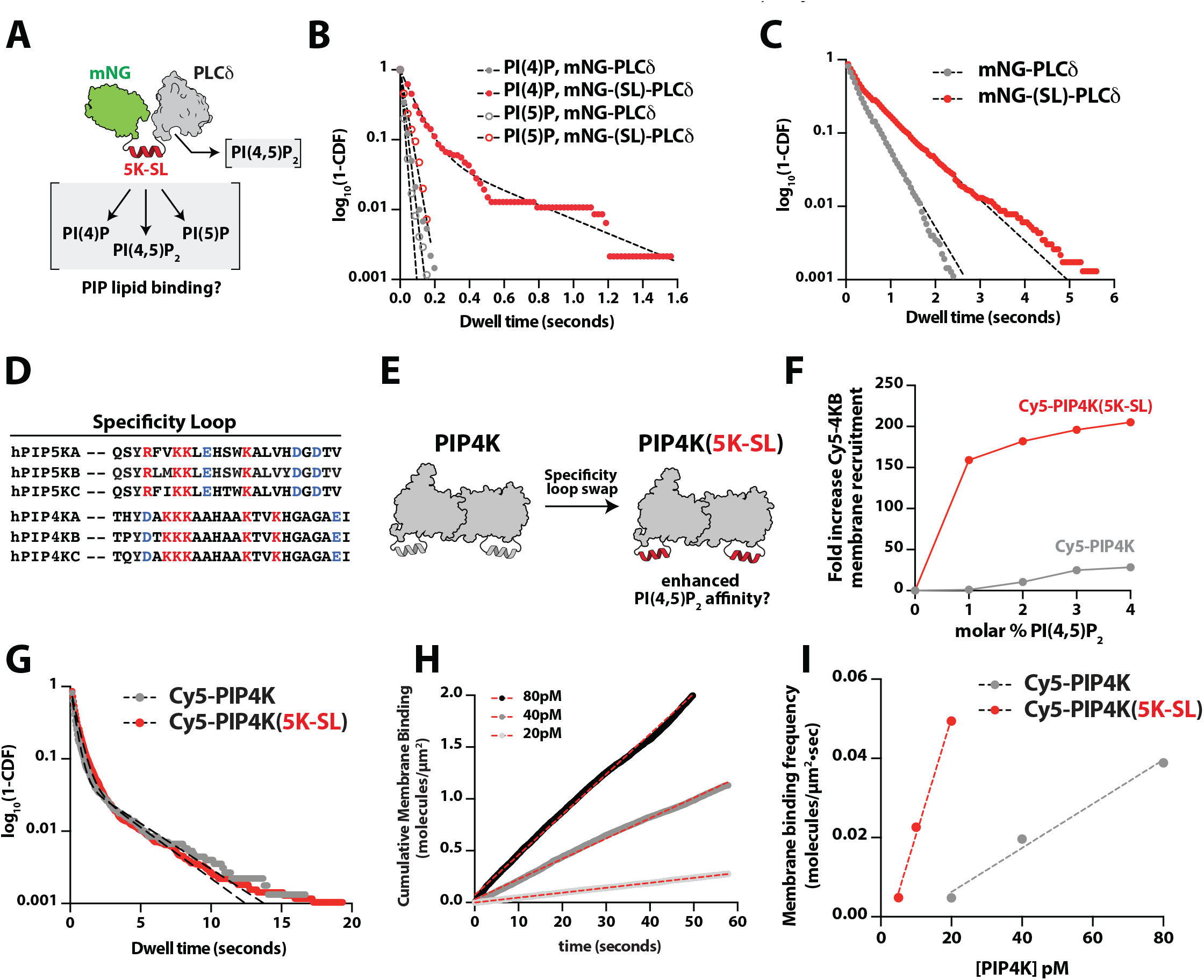
The PIP5K specificity loop promotes PI(4,5)P_2_ lipid binding. **(A)** Cartoon showing the structure of the mNG-PIP5K specificity loop-PLCδ chimeric protein. The expected PIP lipid binding interactions are indicated. **(B)** Single molecule dwell time distributions of mNG-PLCδ and mNG-(5K-SL)-PLCδ measured on membranes containing either 4% PI(4)P or 4% PI(5)P. **(C)** Single molecule dwell time distributions of mNG-PLCδ and mNG-(5K-SL)-PLCδ measured on membranes containing 4% PI(4,5)P_2_. **(D)** Sequence alignment of PIP5K and PIP4K specificity loops. **(E)** Cartoon showing the PIP4K to PIP5K specificity loop swap. **(F)** Quantification of the fold increase in localization of Cy5-PIP4K and Cy5-PIP4K(5K-SL) on membranes containing different molar concentrations of PI(4,5)P_2_. The fold change was calculated relative to the measured membrane density in the absence of PI(4,5)P_2_. **(G)** Single molecule dwell time distributions of Cy5-PIP4K and Cy5-PIP4K(5K-SL) measured on membranes containing 4% PI(4,5)P_2_. **(H)** Plots showing the cumulative membrane binding events measured in the presence of 20-80 pM Cy5-PIP4K. The slope of each curve was calculated by linear regression yielding a binding frequency for each concentration of Cy5-PIP4K. **(I)** Experimental measure of membrane binding frequency (*k*_*ON*_) in the presence of 20-80 pM Cy5-PIP4K or 5-10 pM Cy5-PIP4K(5K-SL).

In addition to regulating substrate recognition, our results indicate that the specificity loop controls PI(4,5)P_2_ lipid binding and contributes to the positive feedback loop observed during PIP5K mediated production of PI(4,5)P_2_. To determine if the PIP5K specificity loop can generally enhance the affinity of other membrane binding proteins we compared the single molecule dwell times of mNG-the common PH-PLCδ domain (**Figure 2C**). Looking at the dwell time distribution of mNG-5K(SL)-PLCδ, we observed a second population of longer dwelling molecules, consistent with the specificity loop enhancing PI(4,5)P_2_ association (**Figure 2C**).

Our recent membrane binding study of the type II PLCδ and mNG-5K(SL)-PLCδ on PI(4,5)P_2_membranes (**Figure 2C**). Both mNG-PLC containing phosphatidylinositol-5-phosphate 4-kinase (PIP4K) revealed that a threshold membrane density of δ and mNG-5K(SL)-PLCδ displayed transient dwell times that were nearly identical (∼150 ms), due to both proteins sharing ∼3% PI(4,5)P_2_ lipids was required to observe robust PIP4K localization on supported lipid bilayers (Wills et al. 2023). By contrast, PIP5K can associate with membranes containing only 1% PI(4,5)P_2_ (Hansen et al. 2019; Wills et al. 2023). To determine whether the difference in PI(4,5)P_2_ dependent membrane association is related to the specific loop sequence variation between PIP4K and PIP5K (**Figure 2D**), we generated a PIP4KB to PIP5KB specificity loop swap referred to as PIP4K(5K-SL) (**Figure 2E**). We quantified the difference in Cy5-PIP4K and Cy5-PIP4K(5K-SL) bulk membrane recruitment on supported membranes containing varying concentrations of PI(4,5)P_2_ (**Figure 2F-2G**). In the presence of 1% PI(4,5)P_2_, Cy5-PIP4K(5K-SL) reached a final equilibrium membrane surface density that was 160-fold greater than Cy5-PIP4K (**Figure 2F**). Visualization of the single molecule membrane binding properties of Cy5-PIP4K and Cy5-PIP4K(5K-SL) revealed that the two kinases had similar dwell times (**Figure 2G**). By contrast, swapping the specificity loop increased the association rate constant (*k*_*ON*_) Cy5-PIP4K(5K-SL) by 5-fold (**Figure 2H-2I**). This is consistent with the role the specificity loop serves in enhancing PIP5K membrane localization to PI(4,5)P_2_ containing membranes. Considering that the PIP4KB to PIP5KB specificity loop swap reduces the isoelectric from 9.7 to 8.4, the observed enhancement in membrane localization for Cy5-PIP4K(5K-SL) cannot be attributed solely to a change in net positive charge. Overall, these data confirm that the PIP5K specificity loop can sense both PI(4)P and PI(4,5)P_2_ lipids. In addition, this data provides one explanation for why PIP4K has a reduced capacity to localize to PI(4,5)P_2_ lipids compared to PIP5K both in vivo and in vitro.

### Molecular basis of the PIP5K high affinity PI(4,5)P_2_ lipid interaction

Having established that the specificity loop serves an important role in regulating the initial membrane docking step of PIP5K, we sought to map the high affinity PI(4,5)P_2_ lipid binding interaction. PIP5K contains numerous basic amino acids on the membrane binding interface that could promote interactions with anionic lipids (Arioka et al. 2004; Fairn et al. 2009). Structural biochemistry studies of zPIP5KA protein crystals soaked with selenate resolved a SeO_4_^2-^molecule bound to basic residues (i.e. K238 and R244) near the ATP binding pocket (Muftuoglu et al. 2016). These residues are part of a conserved sequence, DLKGSxxxR, which is referred to as the PIP binding motif (PIPBM) or substrate binding motif (Muftuoglu et al. 2016) (**Figure 3A**). The PIPBM is considered part of the kinase domain active site and plays an important role in positioning PI(4)P near the ATP γ-phosphate for efficient lipid phosphorylation (Muftuoglu et al. 2016). Consistent with this model, molecular dynamic (MD) simulations have shown that zPIP5KA interacts with PI(4)P lipids through the PIPBM (Amos et al. 2019). Mutation in R244 have been shown to reduce zPIP5KA lipid kinase activity, but paradoxically does not affect the intrinsic ATP hydrolysis rate of zPIP5KA in solution (Muftuoglu et al. 2016).

**Figure 3.**
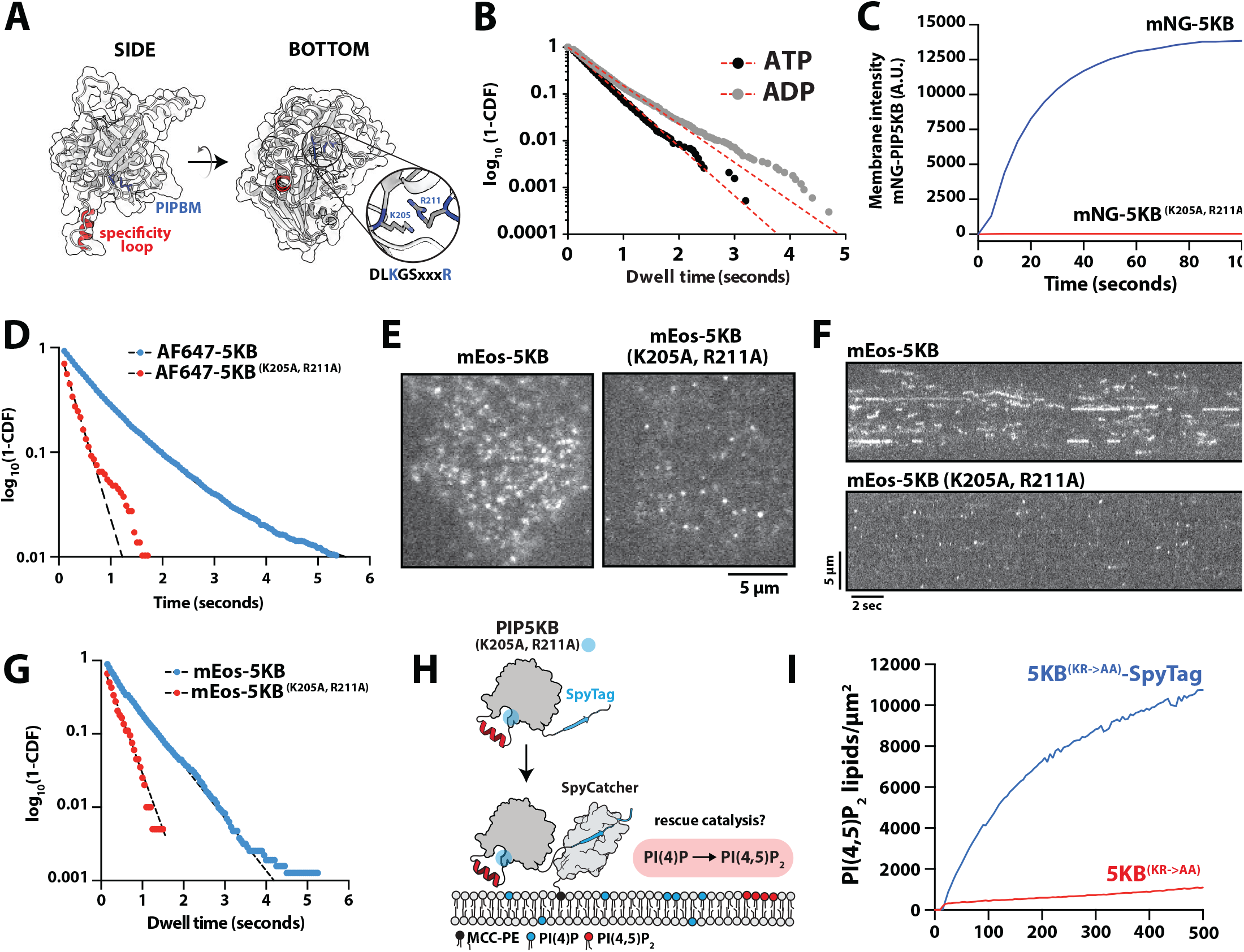
PIP5K engages PI(4,5)P_2_ lipids through the substrate binding motif. **(A)** AlphaFold structure of human PIP5KB validated against zPIP5Kα (5E3U.pdb). Structure highlights the specificity loop (red) and the PIP lipid (or substrate) Binding Motif (PIPBM, blue). **(B)** Single molecule dwell time distributions of AF647-PIP5KB measured in the buffer containing either 1 mM ATP or ADP. **(C)** Bulk membrane recruitment of either 25nM mNG-PIP5KB or mNG-PIP5KB (K205A/ R211A). **(B-C)** Membrane composition: 98% DOPC, 2% PI(4,5)P_2_. **(D)** Single molecule dwell time distribution comparing mNG-PIP5KB and mNG-PIP5KB (K205A/R211A). Membrane composition: 96% DOPC, 4% PI(4,5)P_2_. **(E)** Representative TIRF-M images showing the localization of the mEos3.2-PIP5KB (WT and K205A/R211A) expressed in HEK293T cells. **(F)** Kymograph showing the single molecule plasma membrane binding dynamics of mEos3.2-PIP5KB (WT and K205A/R211A) in HEK293T cells. **(G)** Single molecule dwell time distributions for mEos3.2-PIP5KB (WT and K205A/R211A) measured in HEK293T cells. **(H)** Schematic showing SpyTag-PIP5KB(K205A/ R211A) and SpyCatcher kinase activity assay. **(I)** Kinetics of lipid phosphorylation measured for membrane tethered and non-tethered SpyTag-PIP5KB(K205A/R211A). Production of PI(4,5)P_2_ was monitored in the presence of 20 nM Cy3-PLCδ.

Does the PIPBM regulate PI(4,5)P_2_ binding? Structural modeling of PI(4)P bound to the PIPBM previously suggested that the inositol 5-phosphate of PI(4,5)P_2_ would sterically clash with the γ-phosphate of ATP (Muftuoglu et al. 2016). However, when we measured the single molecule dwell times AF647-PIP5KB diluted in TIRF-M imaging buffer containing either ADP, ATP, or ATPγS, we found that the kinase could associate with PI(4,5)P_2_ membranes independent of the nucleotide chemistry (**Figure 3B**). This suggests that nucleotide binding does not sterically occlude the interaction between PIP5K and PI(4,5)P_2_, which led us to hypothesize that the conserved PIPBM could potentially regulate PI(4,5)P_2_ lipid binding. To test whether the PIPBM is required for the PI(4,5)P_2_ lipid interactions, we introduced mutations that were homologous to zPIP5KA (i.e. K238 and R244) into human PIP5KB (i.e. K205A and R211A). We found that disrupting the PIPBM strongly diminished bulk membrane binding of PIP5K on PI(4,5)P_2_ containing membranes (**Figure 3C**). Consistent with our bulk membrane localization experiment, single molecule characterization of mNG-PIP5KB (K205A, R211A) revealed a 5-fold decrease in the dwell time compared to mNG-PIP5KB (**Figure 3D**). Interestingly, while we did see greatly diminished dwell time, we did not observe a complete loss of PI(4,5)P_2_ binding. This short dwell time could be explained by the remaining specificity loop binding interaction. To determine if our in vitro membrane binding results are consistent in vivo membrane binding dynamics, we expressed mEos3.2-PIP5KB (K205A, R211A) in HEK293T cells and quantified the single molecule plasma membrane dwell time (**Figure 3E**). Compared to mEos3.2-PIP5KB, the PIPBM mutant displayed fewer membrane binding events which can be seen by kymograph analysis (**Figure 3F**). The average dwell time for mEos3.2-PIP5KB (K205A, R211A) was also much shorter compared to mEos3.2-PIP5KB and there were twenty times fewer membrane binding events observed for the same total cellular concentration of fluorescently tagged PIP5KB (**Figure 3G**). This data suggests that the PIPBM serves a critical role in regulating PIP5K plasma membrane localization in cells.

Previous biochemical characterization of the zPIP5KA (R244A) mutant revealed that the PIPBM is critical for phosphorylating PI(4)P to generated PI(4,5) P_2_ (Muftuoglu et al. 2016). Our data also suggests that mutations in the PIPBM dramatically reduce PIP5K membrane binding in the presence of PI(4,5) P_2_. This led us to hypothesize that the previously observed loss in zPIP5KA kinase activity could be due to a disruption in membrane localization. This would be consistent with zPIP5KA (R244A) displaying no change in the ATP hydrolysis rate measured in solution (Muftuoglu et al. 2016). To test this hypothesis, we utilized the SpyTag-SpyCatcher interaction (Li et al. 2014; Zakeri et al. 2012) to rescue PIP5KB (K205A, R211A) membrane localization and (Muftuoglu et al. 2016). Our data also suggests that mutations in the PIPBM dramatically reduce membrane binding of PIP5K. This led us to hypothesize that the previously observed loss in zPIP5KA kinase activity could be due to a disruption in membrane localization. This would be consistent with zPIP5KA (R244A) displaying no change in the ATP hydrolysis rate measured in solution (Muftuoglu et al. 2016). To test this hypothesis, we utilized the SpyTag-SpyCatcher interaction (Li et al. 2014; Zakeri et al. 2012) to rescue PIP5KB (K205A, R211A) membrane localization and lipid kinase activity (**Figure 3H**). For measure lipid kinase activity (**Figure 3H**). For these experiments, SpyCatcher was conjugated to a bilayer via a maleimide lipid. SpyTag-mNG-PIP5KB (K205A, R211A) was then flowed over the supported membrane in buffer lacking ATP and allowed to form an isopeptide bond with membrane conjugated SpyCatcher. After a desired density of SpyTag-mNG-PIP5KB (K205A, R211A) was reached, unbound kinase was washed out of the sample chamber. The kinase reaction was then initiated by adding ATP containing buffer and the PI(4,5)P_2_ biosensor, Cy3-PLCδ. Consistent with the previous characterization of zPIP5KA (R244A), we found that non-membrane tethered SpyTag-PIP5KB (K205A, R211A) mutant was unable to catalyze PI(4,5)P_2_ production in solution (**Figure 3I**). However, upon tethering to SpyTag-PIP5KB (K205A, R211A) to SLBs via SpyCatcher we were able to localize and rescue PI(4,5)P_2_ production. Taken together, these data confirm that the site of high affinity binding and positive feedback come from binding PI(4,5)P_2_ lipids in the PIP lipid binding motif (PIPBM). This high affinity binding interactions controls membrane localization of PIP5K both in vitro and in vivo.

### The PIPBM displays broad specificity for anionic lipids

While the canonical activity of PIP5K is the phosphorylation of PI(4)P to produce PI(4,5)P_2_, PIP5K can also phosphorylate PI(3)P and PI(5)P, albeit less efficiently compared to PI(4)P (Kunz et al. 2002; Muftuoglu et al. 2016). In addition, PIP5K has also been shown to catalyze the synthesis of the important signaling lipid PI(3,4,5)P_3_ using PI(3,4)P_2_ as a substrate (Mufutoglu et al. 2016). Having established that the PIPBM regulates membrane interactions with PI(4,5) P_2_ we aimed to determine whether the PIPBM controls broad specificity for anionic lipids. For this, we created supported lipid bilayers and performed single particle tracking of mNG-PIP5K in the presence of various types of PIP lipids (**Figure 4A**). Overall, we found that mNG-PIP5K bound singly phosphorylated PI(4)P and PI(5)P lipids with a similarly low affinity, while interactions with either doubly or triply phosphorylated PIP lipids (PI(3,4) P_2_, PI(4,5)P_2_, and PI(3,4,5)P_3_) displayed longer dwell times (**Figure 4A**). To test whether PIP5KB prefers to bind doubly and triply phosphorylated PIP lipids due to the structure of the inositol head group versus the total negative charge, we examined the membrane binding properties of mNG-PIP5KB in the presence of PS lipids. We chose phosphatidylserine (PS) because its headgroup has a distinct chemical structure compared to PIP lipids but retains a negative charge. PS lipids have also previously been shown to activate PIP5K, but the molecular basis remains unclear. In agreement with published data, we found that increasing the molar percentage of PS enhanced PIP5K activity, with a physiologically relevant concentrations of 20% PS lipids increasing the kinase activity 42.5-fold compared to membranes that contained 2% PI(4)P and no PS lipids (**Figure 4B**). Correlated with the observed increase in kinase activity, mNG-PIP5KB localization displayed a nonlinear response to PS membrane density. At the physiologically relevant concentration of 20% PS, mNG-PIP5KB membrane localization increased 60-fold compared to membranes lacking PS lipids (**Figure 4C**). Next, we compared the single molecule dwell times of AF647-PIP5KB associated with supported membranes containing either 4% PI(4,5)P_2_ or 20% PS lipids. In the presence of either of these lipid species, AF647-PIP5KB displayed similarly long membrane dwell times (**Figure 4D**). To determine if the interaction with PS lipids was mediated by the PIPBM, we measured the bulk membrane localization of mNG-PIP5K (K205A, R211A). Like our characterization of mNG-PIP5K (K205A, R211A) membrane binding in the presence of PI(4,5)P_2_ (**Figure 3C**), the PIPBM mutant was unable to localize to membranes containing 20% PS lipids (**Figure 4E-4F**). Taken together, these data demonstrate that PIP5K is localized to PS through association of the lipid with the PIPBM, supporting a model in which the PIPBM functions as a broad specificity anionic lipid sensor.

**Figure 4.**
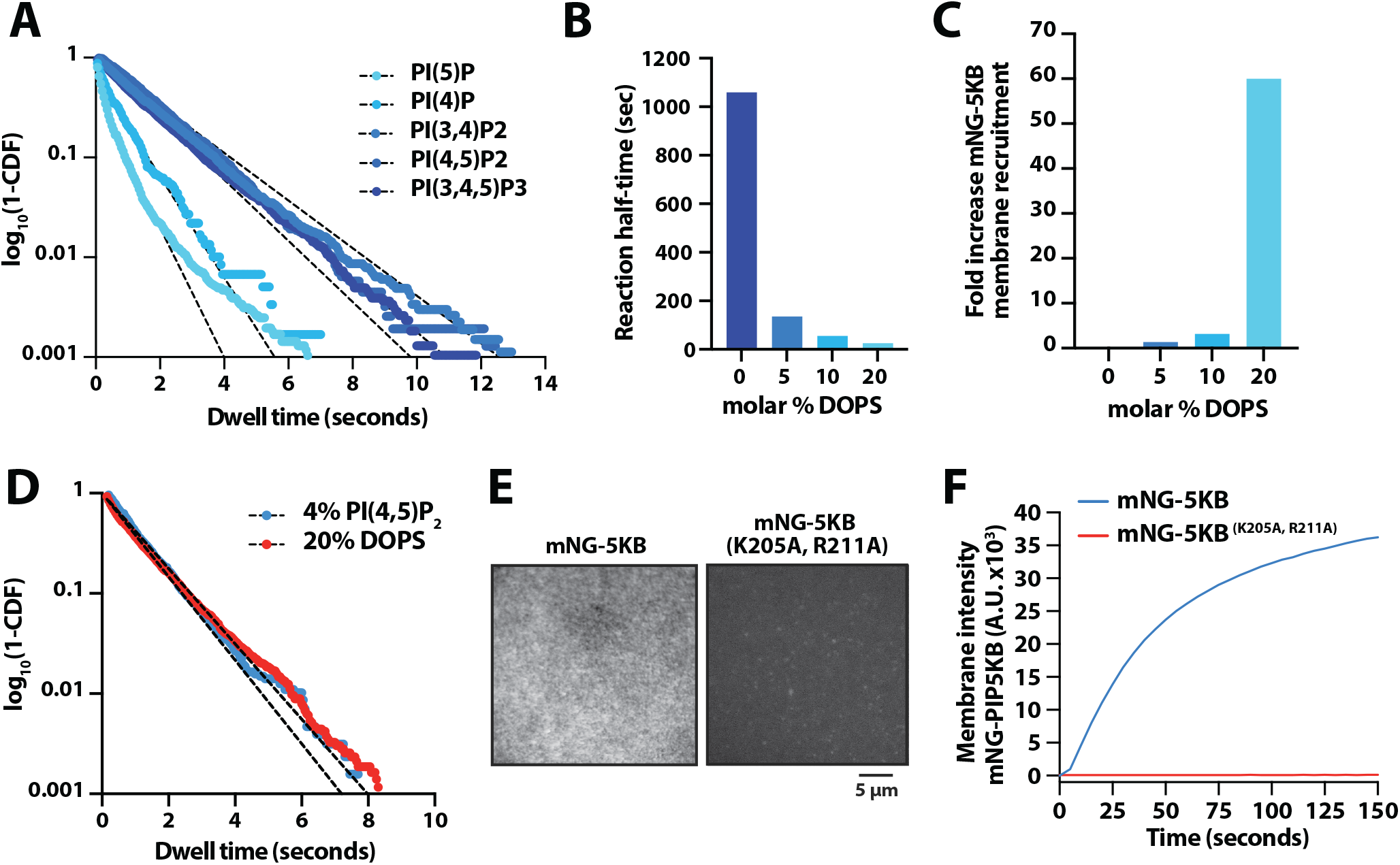
The PIP5K PIPBM exhibits broad specificity for anionic lipids. **(A)** Single molecule dwell time distributions measured in the presence of AF647-PIP5KB on SLBs containing 2% of the indicated PIP lipid. **(B)** Kinetics of PIP5KB dependent lipid phosphorylation measured in the presence of 0-20% DOPS. All membranes contain 2% PI(4)P and reactions were monitored using 20 nM Cy3-PLCδ. Kinetics were quantified by comparing the half-time for reaction completion. **(C)** Quantification of the fold increase of membrane recruitment measured in the presence of 15 nM mNG-PIP5KB on SLBs containing 0-20% DOPS. The fold increase in membrane binding we calculated relative to the membrane intensity of mNG-PIP5K measured in the absence of DOPS. **(D)** Single molecule dwell time analysis of AF647-PIP5KB in the presence of 20% DOPS or 2% PI(4,5)P_2_. **(E)** Representative TIRF-M images showing the membrane localization of 15 nM mNG-PIP5K or mNG-PIP5K (K205A/R211A). Membrane composition: 20% DOPS, 80% DOPC. **(F)** Quantification of bulk recruitment data shown in (E).

### Deciphering the relationship between PIP5K membrane recruitment and catalysis

Our membrane binding studies of the PIPBM mutant revealed that residues that have previously been shown to be critical for lipid kinase activity also regulate PIP5K membrane localization. This raised questions about whether other commonly studied “kinase dead” mutants are defective in membrane binding. For this reason, we aimed to isolate a PIP5K “kinase dead” mutant that lacks lipid phosphorylation activity, but retains the membrane binding dynamics characteristic of the wild type enzyme. To create a separation of function mutant, we rationalized that PIP5K must have a functional PIPBM and specificity loop. Looking at existing structural biochemistry data of zPIP5KA (Muftuoglu et al. 2016), we found that residue D350 helps coordinate a Mg^2+^/Mn^2+^ ions in the active site (**Figure 5A**). Mutations in this residue are predicted to destabilize the transition state following nucleophilic attack of the ATP gamma phosphate. In addition, residues K138 and D266 serve important roles in stabilizing ATP in the active site. Using the mNG-PIP5K single molecule TIRF assay we quantified the membrane binding properties of fluorescently labeled K138A, D266A, and D350A on supported membrane containing 2% PI(4,5)P_2_ (**Figure 5B**). We found only the D266K mutant exhibit membrane binding properties that phenocopied the wild type kinase (**Figure 5C**). This was observed when we measure the single molecule dwell times (**Figure 5C**) and bulk membrane recruitment (**Figure 5D**). Next, we tested the catalytic efficiency of each mutant on supported membranes containing an initial concentration of 4% PI(4)P and no additional anionic lipids. Under these conditions, none of the mutants displayed significant levels of activity compared to the wild type kinase (**Figure 5E**). When we repeated this analysis on supported membranes containing 20% PS lipids, we were able to strongly stimulate the activity of PIP5KB (D350A) through enhanced membrane localization (**Figure 5F**). However, the K138A and D266K mutants still displayed weak activity even though their membrane localization dynamics were most similar to the wild type kinase. Together, these results indicate that some, but not all, reported “kinase dead” mutants are defective in catalysis because of their inability to bind to PI(4,5) P_2_ which is critical for positive feedback.

**Figure 5.**
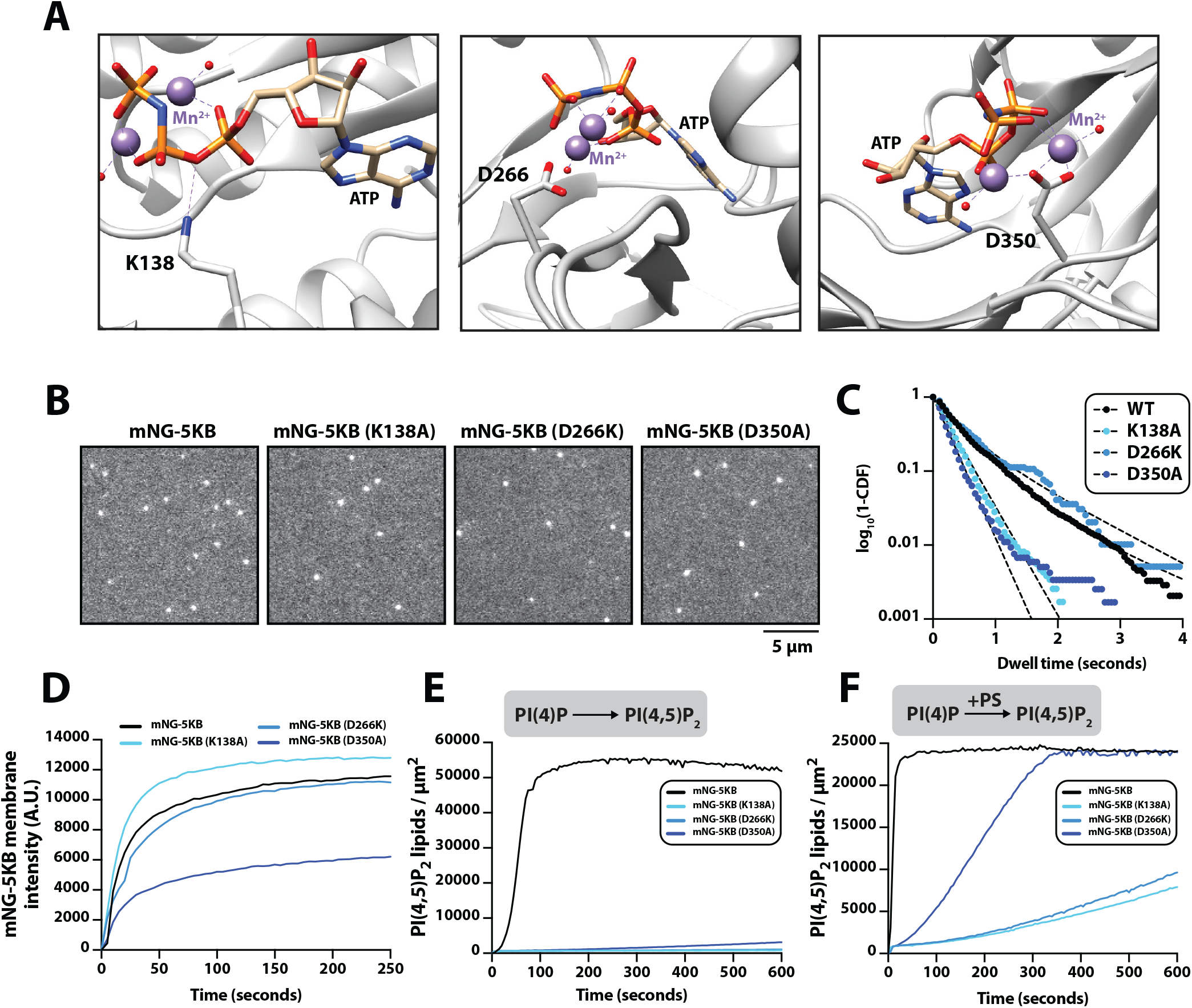
Deciphering the relationship between PIP5K catalysis and membrane recruitment. **(A)** Structure of PIP5K (5E3U.pdb) showing the position of residues that are important for catalysis. **(B)** Representative TIRF-M images showing the localization of mNG-PIP5KB (WT and mutants). **(C)** Single molecule dwell time distributions for mNG-PIP5KB (WT and mutants) measured. **(B-C)** Membrane composition: 98% DOPC, 2% PI(4,5)P_2_. **(D)** Bulk membrane recruitment measured by TIRF-M in the presence of 20 nM mNG-PIP5KB (WT and mutants). Membrane composition: 98% DOPC, 2% PI(4,5)P_2_. **(E-F)** Lipid kinase activity measured using TIRF-M in the presence of 10 nM PIP5KB (WT and mutants). Production of PI(4,5)P_2_ was monitored in the presence of 20 nM Cy3-PLCδ. **(E)** Membrane composition: 96% DOPC, 4% PI(4)P. **(F)** Membrane composition: 78% DOPC, 2% PI(4)P, 20% DOPS.

## DISCUSSION

Class I PIP kinases – PIP5Kα, β, and γ – are critical regulators of cellular homeostasis. Recent work has identified PIP5K as a potential cancer therapeutic target. To improve the efficacy of PIP5K inhibitors, deciphering the mechanisms that regulate lipid specificity and membrane localization of PIP5K is paramount. Currently, the sequence of molecular interactions that allow PIP5K to associate with membranes containing anionic lipids remains unclear. Previous structural biochemistry studies have implicated both the specificity loop and PIP lipid binding motif (PIPBM) of PIP5K in regulating lipid specificity and kinase activity (Liu et al. 2016; Muftuoglu et al. 2016). While these structural studies established a foundational model for PIP5K function, measurements with higher spatial and temporal resolution are needed to directly visualize and determine the role the specificity loop and PIPBM serve in controlling PIP5K localization and activity on membranes. Using smTIRF microscopy and mutational analysis we determined that PIP5K associates with anionic lipids through a two-step mechanism that robustly docks the kinase on membranes. The initial membrane docking step utilizes the specificity loop to engage either PI(4) P or PI(4,5)P_2_ lipids. This is consistent with molecular dynamic simulations that reported this potential mechanism for controlling membrane binding of PIP5K (Amos et al. 2019). We find that mutations that disrupt the alpha helical conformation of the specificity loop drastically reduced the association rate (*k*_*ON*_) and partially increased the dissociation rate (*k*_*OFF*_) of PIP5K. Similarly, we found that the PIP5KB specificity loop mutant, W365P, displayed a reduced plasma membrane binding frequency in cells. However, the single molecule dwell times of W365P were comparable to wild type PIP5KB in vivo. Overall, the specificity loop is dispensable for stable membrane association. However, the decrease in *k*_*ON*_ suggests the specificity loop helps orient the kinase so that the PIPBM can more efficiently engage anionic lipids.

PIP5Ks specificity loop has been shown to be critical for substrate specificity and catalysis (Kunz et al. 2000, 2002; Liu et al. 2016). However, the exact mechanism by which this motif gives rise to lipid specificity remains unclear. In some cases, the specificity loop has been described as a broad membrane lipid sensor. In other cases, the specificity loop has been shown to strictly engage PI(4)P. Here, we report that the specificity loop confers binding to both PI(4)P and PI(4,5)P_2_ lipids. We found that a chimeric protein, mNG-(5K-SL)-PLCδ, containing the specificity loop derived from PIP5K displayed a gain of function in ability to localize to PI(4) P and PI(4,5)P_2_ containing membranes. By contrast, mNG-(5K-SL)-PLCδ did not exhibit any specificity for PI(5)P. The ability of the PIP5K specificity loop to bind PI(4,5)P_2_ was also confirmed by attaching this sequence to PIP4K, the kinase that typically uses PI(5)P as a substrate. This resulted in a dramatic enhancement in the ability of PIP4K to bind PI(4,5)P_2_ lipids, further supporting the specificity loop contribute to PI(4,5)P_2_-mediated membrane recruitment and positive feedback during catalysis. Together, these findings provide the first mechanistic model for how the specificity loop selectively binds to PI(4)P and PI(4,5)P_2_ lipids. Our results support a model in which the specificity loop engages both the substrate and product.

Previous studies have described the PIP binding motif (PIPBM) as the active site of the kinase, which controls specificity for PI(4)P (Muftuoglu et al. 2016). However, we found that the PIPBM also displays a high affinity for the lipid product, PI(4,5)P_2_. Through mutational analysis of the PIPBM, we identified the residues required for the high affinity interaction between PIP5K and PI(4,5)P_2_ lipids. Mutations in the PIPBM dramatically reduced the bulk membrane binding of PIP5K, and greatly reduced the single molecule dwell times measured by TIRF-M. When we measured the lipid specificity of the PIPBM, we found it could broadly associate with anionic lipids. Consistent with this model, a PIP5K mutant lacking the PIPBM motif did not productively localize to the plasma membrane in vivo. In vitro, PIP5K bound weakly to singly phosphorylated PI(4)P and PI(5)P lipids but displayed progressively higher affinities for PI(4,5)P_2_, PI(3,4)P_2_, and PI(3,4,5)P_3_. PIP5K could also strongly associate with containing 20% PS lipids. Under these conditions, PIP5K membrane recruitment mirrored what we observed in the presence of 2% PIP lipids. Note that while our experiments utilized PIP5KB for biochemical analysis, both PIP5KA and PIP5KC contain homologous specificity loops and PIPBMs that are expected to function in an identical manner.

Our detailed biochemical analysis of PIP5K membrane targeting and lipid specificity provide novel insights into how PIP5K localizes to membranes, which has important implications for how PIP5K is activated by negatively charged lipids like phosphatidic acid and phosphatidylserine (Cockcroft 2009; Ishihara et al. 1998; Jenkins, Fisette, and Anderson 1994). In vivo studies of PIP5K localization have always shown that each of the PIP5K isoforms localize primarily to the plasma membrane. Despite this, no study has been able to fully explain this localization pattern. Until now, the field has hypothesized that PIP5K localizes to the plasma membrane through interactions with it’s substrate, PI(4)P. However, PI(4)P also exists on Golgi membranes but does not strongly localize PIP5K to that organelle. PIP5Ks product exists at the plasma membrane, and our work demonstrates that PIP5K have a higher affinity to this lipid than to PI(4)P, which likely plays a role in controlling PIP5K’s localization. However, studies that perturbed PI(4,5)P_2_ at the plasma membrane did not see complete disruption of PIP5Ks localization, suggesting that PI(4,5)P_2_ is not the sole factor driving localization. Our findings of high affinity binding to physiologic concentrations of PS lipids suggests an alternative mechanism for controlling PIP5K localization to the plasma membrane. While PS lipids are present at most intracellular membranes, its only at the plasma membrane that PS lipids reach a molar density of 20%. Testing the hypothesis that PS lipids drive localization of PIP5K to the plasma membrane is currently challenging, because we lack methods to acutely eliminate PS lipids without having pleiotropic effects. Further cell biological studies may be able to elucidate whether either or both mechanisms are leveraged by cells to recruit and activate PIP5K during signal transduction and response.

## AUTHOR CONTRIBUTIONS

Resources: B.R.D., K.A.F., and S.D.H.

Experiments and investigation: B.R.D., K.A.F., and S.D.H.

Data Analysis: B.R.D., K.A.F., and S.D.H.

Conceptualization: B.R.D., K.A.F., and S.D.H.

Interpretation: B.R.D., K.A.F., and S.D.H.

Data curation: B.R.D. and S.D.H.

Writing – Review and editing: B.R.D., K.A.F., and S.D.H.

Writing – Original draft: B.R.D. and S.D.H.

Supervision: S.D.H.

Project administration: S.D.H.

Funding acquisition: S.D.H.

## RESOURCE AVAILABILITY

All the information needed for interpretation of the data is presented in the manuscript or the supplemental material. Plasmids related to this work are available upon request.

## MATERIALS & METHODS

### Molecular Biology

The gene coding the PH domain derived from human phospholipase C-δ1 (PLCδ Accession #P51178.2), human phosphatidylinositol 4-phosphate 5-kinase type-1 beta (hPIP5KB; Uniprot #O14986) and human phosphatidylinositol 5-phosphate 4-kinase type-2 beta (hPIP4KB; Uniprot #P78356) were derived from codon optimized genes synthesized by GeneArt (Invitrogen). Gene sequences were subcloned into either bacterial, insect cell, or mammalian expression vectors using Gibson assembly (Gibson et al. 2009). Plasmids containing PIP5KB mutations (i.e. W365P, D51R, K205A, R211A, K138A, D350A, or D266K) were made through site-directed mutagenesis using the PfuUltra High-Fidelity DNA polymerase (Agilent, cat# 600380). Chimeric mNG-PIP5K, mNG-PLCδ, and mNG-Specificity Loop-PLCδ were created using PCR amplification of genes via AccuPrime *Pfx* Master Mix (ThermoFisher, Cat#12344040) then combined with a digested plasmid using Gibson Assembly. The complete open reading frame of all vectors used in this study were sequenced by Azenta (formerly Genewiz) and Plasmidsaurus (University of Oregon) to ensure constructs lacked deleterious mutations. Each protein expression construct was screened for optimal yield and solubility in either bacteria (BL21 DE3 Star, Rosetta, etc.) or *Spodoptera frugiperda* (Sf9) insect cells. See Supplementary file for the amino acid sequences used during recombinant protein expression or transient transfection in HEK293T cells. Note that there have previously been inconsistencies in nomenclature between human and mouse PIP5K paralogs. In this manuscript, PIP5Kβ refers to the human PIP5Kβ paralog. All plasmids created and used within this manuscript function under this human nomenclature.

### Protein purification

#### PIP5KB and mNG-PIP5K

Gene sequences encoding PIP5KB and mutants (i.e. W365P, D51R, and K205A/ R211A) were cloned into FastBac1 vectors in frame with a N-terminal his_6_-MBP-TEV-GGGGG or his_6_-TEV-mNG-GGGGG, and expressed under the polyhedrin (pH) promoter. BACMIDS and baculoviruses were generated as previously described (Hansen et al. 2019). For large protein expression, high five cells were infected with baculovirus using an optimized multiplicity of infection (MOI), typically 2% vol/vol. Infected insect cells were grown for 48 hours at 27^0^C in ESF 921 Serum-Free Insect Cell Culture medium (Expression Systems, Cat# 96-001-01). Cells were then harvested by centrifugation, washed with 1x PBS [pH 7.2], resuspended in cell storage buffer (1x PBS [pH 7.2], 10% glycerol, 2x Sigma protease inhibitor table), and then stored in the -80ºC freezer. For purification, frozen insect cell pellets for 2-4 liters of liquid culture were thawed at room temperature in a water bath and lysed into buffer containing 50 mM Na_2_HPO_4_ [pH 8.0], 10 mM imidazole, 400 mM NaCl, 1 mM PMSF (added twice, once before homogenization and once after), 5 mM BME, 100 μg/mL DNase, SIGMAFAST protease inhibitor cocktail tablets, EDTA-free (Sigma, cat# S8830-20TAB) per 100 mL lysis buffer. Cells in this buffer were lysed using a dounce homogenizer. Lysate was clarified by centrifugation at 36,000 rpm (140,000 x *g*) for 60 minutes under vacuum using a Beckman Ti-45 rotor at 4ºC. Lysate was then batch bound to 5 mL of Ni-NTA Agarose (Qiagen, Cat# 30230) resin at 4ºC for 2 hours in a beaker set on a stir plate. Resin was then collected in 50 mL tubes, centrifuged, and washed with buffer containing 50 mM Na_2_HPO_4_ [pH 8.0], 10 mM imidazole, 400 mM NaCl, and 5 mM BME and centrifuged again before being transferred to gravity flow column in more wash buffer. Ni-NTA resin with (His)_6_-MBP-(Asn)_10_-TEV-GGGGG-PIP5K bound was then eluted into buffer containing 500 mM imidazole. Peak fractions were pooled, combined with 200 μg/mL his6-TEV(S291V) protease, and dialyzed against 4 liters of buffer containing 20 mM Tris [pH 8.0], 200 mM NaCl, 2.5 mM BME for 16-18 hours at 4ºC. Dialysate was then combined 1:1 with 20 mM Tris [pH 8.0], 1 mM TCEP (∼100 mM NaCl final). Precipitation was removed by centrifugation and 0.22 μm syringe filtration. Clarified dialysate was then bound to a MonoS cation exchange column (GE Healthcare, Cat# 17-5168-01) equilibrated in 20 mM Tris [pH 8.0], 100 mM NaCl, 1 mM TCEP buffer. Proteins were resolved over a 10-100% linear gradient (0.1-1 M NaCl, 45 CV, 45 mL total, 1 mL/min flow rate). PIP5K homologs and paralogs typically eluted from the MonoS column in the presence of 370-450 mM NaCl. Peak fractions containing PIP5K were pooled, concentrated in a 30 kDa MWCO Vivaspin 6 centrifuge tube (GE Healthcare, Cat# 28-9323-17), and loaded onto a 24 mL Superdex 200 10/300 GL (GE Healthcare, Cat# 17-5174-01) size exclusion column equilibrated in 20 mM Tris [pH 8.0], 200 mM NaCl, 10% glycerol, 1 mM TCEP. Peak fractions were concentrated in a 30 kDa MWCO Vivaspin 6 centrifuge tube and snap frozen at a final concentration of 10-40 μM using liquid nitrogen. GGGGG-PIP5KB were labeled with Alexa647-LPETGG using sortase mediated peptide ligation as previously described (Hansen et al. 2019).

#### PIP4K2B

Codon optimized gene sequence encoding human PIP4K2B isoform 2 (Uniprot # P78356) was cloned into a pETM derived bacterial expression vector to create the following fusion protein: his_6_-SUMO3-GGGGG-PIP4K2B (1-416aa). Recombinant PIP4K2B was expressed in BL21(DE3) Star *E. coli* as previously described (Wills et al. 2022). Using 2-4 liters of Terrific Broth, bacterial cultures were grown at 37ºC until OD_600_=0.6. Cultures were then shifted to 18ºC for 1 hour to cool down. Protein expression was induced with 50 μM IPTG and bacteria expressed protein for 20 hours at 18ºC before being harvested by centrifugation. For purification, cells were lysed into buffer containing 50 mM Na_2_HPO_4_ [pH 8.0], 400 mM NaCl, 0.4 mM BME, 1 mM PMSF (add twice, 15 minutes intervals), DNase, 1 mg/mL lysozyme using a microtip sonicator. Lysate was centrifuged at 16,000 rpm (35,172 x *g*) for 60 minutes in a Beckman JA-17 rotor chilled to 4ºC. Lysate was circulated over 5 mL HiTrap Chelating column (GE Healthcare, Cat# 17-0409-01) that had been equilibrated with 100 mM CoCl_2_ for 1 hour, washed with MilliQ water, and followed by buffer containing 50 mM Na_2_HPO_4_ [pH 8.0], 400 mM NaCl, 0.4 mM BME. Recombinant PIP4K2B was eluted with a linear gradient of imidazole (0-500 mM, 8 CV, 40 mL total, 2 mL/min flow rate). Peak fractions were pooled, combined with 50 μg/mL of his6-SenP2 (SUMO protease), and dialyzed against 4 liters of buffer containing 25 mM Na_2_HPO_4_ [pH 8.0], 400 mM NaCl, and 0.4 mM BME for 16-18 hours at 4ºC. Following overnight cleavage of the SUMO3 tag, dialysate containing his6-SUMO3, his6-SenP2, and GGGGG-PIP4K2B was recirculated for at least 1 hr over a 5 mL HiTrap(Co^+2^) chelating column. Flow-through containing GGGGG-PIP4K2B was then concentrated in a 30 kDa MWCO Vivaspin 6 before loading onto a Superdex 200 size exclusion column equilibrated in 20 mM HEPES [pH 7], 200 mM NaCl, 10% glycerol, 1 mM TCEP. In some cases, cation exchange chromatography was used to increase the purity of GGGGG-PIP4K2B before loading on the Superdex 200. In those cases, we equilibrated a MonoS column 20 mM HEPES [pH 7], 100 mM NaCl, 1 mM TCEP buffer. PIP4K2B (pI = 6.9) bound to the MonoS was resolved over a 10-100% linear gradient (0.1-1 M NaCl, 30 CV, 30 mL total, 1.5 mL/min flow rate). Peak fractions collected from the Superdex 200were concentrated in a 30 kDa MWCO Vivaspin 6 centrifuge tube and snap frozen at a final concentration of 20-80 μM using liquid nitrogen.

#### PIP4K2B Specificity Loop Swap (SLS)

Codon optimized gene sequence encoding human PIP4K2B isoform 2 (Uniprot # P78356) was modified by site directed mutagenesis to swap the PIP4K2B specificity loop (372-384aa) with PIP5KB specificity loop (353-373aa) we refer to as PIP4K2B SLS. This construct was cloned into a pETM derived bacterial expression vector to create the following fusion protein: his6-SUMO3-GGGGG-PIP4K2B(5K-SL). Recombinant PIP4K2B was expressed in BL21(DE3) Star *E. coli*. In 2 liters of Terrific Broth, bacterial cultures were grown at 37ºC until OD_600_=0.6. Cultures were then shifted to 18ºC for 1 hour to cool down. Protein expression was induced with 50 μM IPTG and bacteria expressed protein for 20 hours at 18ºC before being harvested by centrifugation. For purification, cells were lysed into buffer containing 50 mM Na_2_HPO_4_ [pH 8.0], 400 mM NaCl, 0.4 mM BME, 1 mM PMSF, DNase, 1 mg/mL lysozyme using a microtip sonicator. Lysate was centrifuged at 16,000 rpm (35,172 x *g*) for 60 minutes in a Beckman JA-17 rotor chilled to 4ºC. Lysate was circulated over 5 mL HiTrap Chelating column (GE Healthcare, Cat# 17-0409-01) that had been equilibrated with 100 mM CoCl_2_ for 1 hour, washed with MilliQ water, and followed by buffer containing 50 mM Na_2_HPO_4_ [pH 8.0], 400 mM NaCl, 0.4 mM BME. Recombinant PIP4K2B was eluted with a linear gradient of imidazole (0-500 mM, 8 CV, 40 mL total, 2 mL/min flow rate). Peak fractions were pooled, combined with 50 μg/mL of his6-SenP2 (SUMO protease), and dialyzed against 4 liters of buffer containing 25 mM Na_2_HPO_4_ [pH 8.0], 400 mM NaCl, and 0.4 mM BME for 16-18 hours at 4ºC. Following overnight cleavage of the SUMO3 tag, dialysate containing his6-SUMO3, his6-SenP2, and GGGGG-PIP4K2B was recirculated for at least 1 hr over a 5 mL HiTrap(Co^+2^) chelating column. Flow-through containing GGGGG-PIP4K2B was then concentrated in a 30 kDa MWCO Vivaspin 6 before loading onto a Superdex 200 size exclusion column equilibrated in 20 mM HEPES [pH 7], 200 mM NaCl, 10% glycerol, 1 mM TCEP. In some cases, cation exchange chromatography was used to increase the purity of GGGGG-PIP4K2B before loading on the Superdex 200. In those cases, we equilibrated a MonoS column 20 mM HEPES [pH 7], 100 mM NaCl, 1 mM TCEP buffer. PIP4K2B (pI = 6.9) bound to the MonoS was resolved over a 10-100% linear gradient (0.1-1 M NaCl, 30 CV, 30 mL total, 1.5 mL/min flow rate). Peak fractions collected from the Superdex 200 were concentrated in a 30 kDa MWCO Vivaspin 6 centrifuge tube and snap frozen at a final concentration of 20-80 μM using liquid nitrogen.

#### PLCδ-PH domain

This protein was expressed and purified as previously described (Hansen et al. 2019). Briefly, human PLCδ-PH domain (11-140aa) was expressed in BL21 (DE3) Star *E. coli* as a his_6_-SUMO3-(Gly)_5_-PLCδ (11-140aa) fusion protein. Following growth at 37ºC in Terrific Broth to an OD_600_ of 0.8, cultures were shifted to 18ºC for 1 hour, induced with 0.1 mM IPTG, and allowed to express protein for 20 hours at 18ºC before being harvested. Cells were lysed into 50 mM Na_2_HPO_4_ [pH 8.0], 300 mM NaCl, 0.4 mM BME, 1 mM PMSF, 100 μg/ mL Dnase using a microfluidizer. Lysate was then centrifuged at 16,000 rpm (35,172 x *g*) for 60 minutes in a Beckman JA-17 rotor chilled to 4ºC. Lysate was circulated over 5 mL HiTrap Chelating column (GE Healthcare, Cat# 17-0409-01) charged with 100 mM CoCl_2_ for 1 hour. Bound protein was then eluted with a linear gradient of imidazole (0-500 mM, 8 CV, 40 mL total, 2 mL/min flow rate). Peak fractions were pooled, combined with SUMO protease (50 μg/mL final concentration), and dialyzed against 4 liters of buffer containing 50 mM Na_2_HPO_4_ [pH 8.0], 300 mM NaCl, and 0.4 mM BME for 16-18 hours at 4ºC. Dialysate containing SUMO cleaved protein was recirculated for 1 hr over a 5 mL HiTrap Chelating column. Flow-through containing (Gly)_5_-PLCδ (11-140aa) was then concentrated in a 5 kDa MWCO Vivaspin 20 before being loaded on a Superdex 75 size exclusion column equilibrated in 20 mM Tris [pH 8.0], 200 mM NaCl, 10% glycerol, 1 mM TCEP. Peak fractions containing (Gly)_5_-PLCδ (11-140aa) were pooled and concentrated to a maximum concentration of 75 μM (1.2 mg/mL) before snap freezing with liquid nitrogen and storage at -80ºC.

#### mNG-PLCδ-PH and mNG-(5K-SL)-PLCδ-PH

Genes encoding chimeric mNeonGreen-hPLCδ-PH domain (11-140aa) and mNeonGreen-hPIP5K Specificity Loop (353-373)-hPLCδ-PH domain (11-140aa) were cloned into bacterial expression vectors. Each construct was expressed in BL21 (DE3) Star *E. coli* as his_10_-TEV fusion proteins. Bacteria were grown at 37ºC in Terrific Broth to an OD_600_ of 0.8. Cultures were induced with 0.1 mM IPTG and allowed to express protein for 20 hours at 18ºC before being harvested. Cells were lysed into 50 mM Na_2_HPO_4_ [pH 8.0], 400 mM NaCl, 0.5 mM BME, 1 mM PMSF, 100 μg/mL DNase using tip sonication (45% amplitude, 5 seconds on, 10 seconds off). Lysate was then centrifuged at 16,000 rpm (35,172 x *g*) for 60 minutes in a Beckman JA-17 rotor chilled to 4ºC. Lysate was circulated over 5 mL HiTrap Chelating column (GE Healthcare, Cat# 17-0409-01) charged with 100 mM CoCl_2_ for 1 hour. Bound protein was then eluted with a linear gradient of imidazole (0-500 mM, 8 CV, 40 mL total, 2 mL/min flow rate). Peak fractions were pooled and combined with 200 μg/mL his6-TEV(S291V) protease and dialyzed against 4 liters of buffer containing 50 mM Na_2_HPO_4_ [pH 8.0], 400 mM NaCl, and 0.4 mM BME for 16-18 hours at 4ºC. Dialysate containing cleaved His_10_-TEV protein was recirculated for 1 hr over a 5 mL HiTrap Chelating column. Flow-through containing mNG-PLCδ or mNG-(5K-SL)-PLCδ were then concentrated in a 50 kDa MWCO Vivaspin 20 before being loaded on a Superdex 75 size exclusion column equilibrated in 25 mM Tris [pH 8.0], 150 mM NaCl, 10% glycerol, 1 mM TCEP. Peak fractions containing either mNG-PLCδ or mNG-(5K-SL)-PLCδ were pooled and concentrated to 4-20μM before snap freezing with liquid nitrogen and storage at -80ºC.

#### Preparation of small unilamellar vesicles

The following lipids were used to generated small unilamellar vesicles (SUVs): 1,2-dioleoyl-sn-glycero-3-phosphocholine (18:1 DOPC, Avanti # 850375C), L-α-phosphatidylinositol-4-phosphate (Brain PI(4)P, Avanti Cat# 840045X), 1,2-dioleoyl-sn-glycero-3-phospho-(1’-myo-inositol-5’-phosphate) (PI(5)P, Avanti Cat# 850152P), L-α-phosphatidylinositol-4,5-bisphosphate (Brain PI(4,5)P_2_, Avanti # 840046X), D-myo-phosphatidylinositol 3,4,5-trisphosphate (PI(3,4,5)P_3_ diC16, Echelon P-3916-100ug), D-myo-phosphatidylinositol 3,4-bisphosphate (PI(3,4)P_2_ diC16, Echelon P-3416-100ug), 1,2-dioleoyl-*sn-*glycero-3-phospho-L-serine (18:1 DOPS, Avanti # 840035C), 1,2-dioleoyl-sn-glycero-3-phosphoethanolamine-N-[4-(p-maleimidomethyl)cyclohexane-carboxamide] (18:1 MCC-PE, Avanti #780201C). To make liposomes, 2 μmoles total lipids are combined in a 35 mL glass round bottom flask with 2 mL of chloroform. Lipids are dried to a thin film using rotary evaporation with the glass round-bottom flask submerged in a 42ºC water bath. The lipid film was then resuspended in 2 mL of PBS [pH 7.2], getting a final concentration of 1 mM total lipids. All lipid mixtures expressed as percentages (e.g. 98% DOPC, 2% PI(4)P) are equivalent to molar fractions. To generate 30-50 nm SUVs, 1 mM total lipid mixtures were extruded through a 0.05 μm pore size 19 mm polycarbonate membrane (Avanti #610002) with filter supports (Avanti #610014) on both sides of the polycarbonate membrane. Extruding hydrated lipids a total of 11 times achieved the desired SUV size.

#### Preparation of supported lipid bilayers

Supported lipid bilayers are formed on 25x75 mm coverglass (IBIDI, #10812). Coverglass first is cleaned with 2% Hellmanex III (Fisher, Cat#14-385-864) heated to 60-70ºC in a glass coplin jar. This was incubated for at least 30 minutes. Once incubated, the coverglass was washed thoroughly with MilliQ water. Once cleaned, the coverglass was then etched in Pirahna solution (1:3, hydrogen peroxide:sulfuric acid) for 10-15 minutes the same day SLBs were formed. Once etched coverglass was again thoroughly rinsed with MilliQ water before being rapidly dried with nitrogen glass. Once dried, glass was adhered to a 6-well sticky-side chamber (IBIDI, Cat# 80608). SLBs were formed by flowing 30 nm SUVs diluted in PBS [pH 7.2] to a total lipid concentration of 0.25 mM and incubated for 30 minutes. IBIDI chambers were then washed with 5 mL of PBS [pH 7.2] to remove non-absorbed SUVs. Membrane defects are blocked for 15 minutes with a 1 mg/mL beta casein (ThermoFisher, Cat# 37528) diluted in 1x PBS [pH 7.4]. Before use as a blocking protein, frozen 10 mg/mL beta casein stocks were thawed, centrifuged for 30 minutes at 21370 x *g*, and 0.22 μm syringe filtered. After blocking SLBs with beta casein, membranes were washed again with 1mL of PBS, followed by 1 mL of kinase buffer before TIRF-M.

#### Membrane conjugation of SpyCatcher

Following beta-casein blocking of SLBs, membranes that contained MCC-PE lipids were washed into 1x PBS [pH 7.2] containing 0.1 mM TCEP. MCC-PE lipids were used to covalently couple SpyCatcher protein onto supported bilayers. For these SLBs, 100 μL of 30 μM SpyCatcher diluted in a 1x PBS [pH 7.2] and 0.1 mM TCEP buffer was added to the IBIDI chamber and incubated for 2 hours at 23ºC. Once the coupling period passed, SLBs with MCC-PE lipids were then washed with 2 mL of 1x PBS [pH 7.2] containing 5 mM beta-mercaptoethanol (BME) and incubated in this buffer for 15 minutes to quench the unreacted maleimide headgroups. SLBs were then washed with 2 mL of 1x PBS to remove unbound protein. Membrane were stored for up to 2 hours in 1x PBS before being washed into imaging buffer to initiate TIRF-M measurements.

#### Kinetics measurements of PI(4,5)P_2_ lipid phosphorylation

The kinetics of PI(4)P phosphorylation were measured on SLBs formed in IBIDI chambers and visualized using TIRF microscopy as previously described (Hansen et al. 2019, 2022). Reaction buffer contained 20 mM HEPES [pH 7.0], 150 mM NaCl, 1 mM ATP, 5 mM MgCl_2_, 0.5 mM EGTA, 20 mM glucose, 200 μg/mL beta casein (ThermoScientific, Cat# 37528), 20 mM BME, 320 μg/mL glucose oxidase (Serva, #22780.01 *Aspergillus niger*), 50 μg/mL catalase (Sigma, #C40-100MG Bovine Liver), and 2 mM Trolox (UV treated, see methods below). Perishable reagents (i.e. glucose oxidase, catalase, and Trolox) were added 5-10 minutes before image acquisition. For all experiments, we monitored the change in PI(4)P or PI(4,5)P_2_ membrane density using a solution concentration of 20 nM Cy3-PLCδ. Density of PIP lipids (lipids/μm^2^) was calculated assuming a footprint of 0.72 nm^2^ for DOPC lipids (Galush, Nye, and Groves 2008; Vacklin, Tiberg, and Thomas 2005).

#### Microscope hardware and imaging acquisition

Single molecule imaging experiments were performed on an inverted Nikon Ti2 microscope using a 100x Nikon objective (1.49 NA) oil immersion TIRF objective. The x-axis and y-axis positions were manually controlled using a Nikon motorized stage and joystick. All images were acquired using a iXion Life 897 EMCCD camera (Andor Technology Ltd., UK). Fluorescently labeled proteins were excited with either a 488 nm, 561 nm, or 637 nm diode laser (OBIS laser diode, Coherent Inc. Santa Clara, CA) controlled by a Vortran laser drive with acousto-optic tunable filters (AOTF) control. The power output measured through the objective for single particle imaging was 1-3 mW. For dual color imaging of mNG-PIP5K localization during Cy3-PLCδ monitored PI(4,5)P_2_ synthesis, samples were excited with 1 mW 488 nm and 1 mW 561 nm light, as measured through the objective. Excitation light was passed through the following dichroic filter cubes before illuminating the sample: (1) ZT488/647rpc and (2) ZT561rdc (ET575LP) (Semrock). Fluorescence emission was detected on an ANDOR EMCCD camera position after a Sutter emission filter wheel housing the following emission filters: ET525/50M, ET600/50M, ET700/75M (Semrock). All experiments were performed at room temperature (23ºC). Microscope hardware was controlled using Nikon NIS elements.

#### Single particle tracking

Fluorescent protein detection and tracking was performed using the ImageJ/Fiji TrackMate plugin (Jaqaman et al. 2008). Data in the form of .nd2 files were loaded in ImageJ. Before being analyzed using TrackMate, data brightness was adjusted for molecules to be easily identifiable. TrackMate was then used to identify and track molecular tracks in these steps: Particles were first identified using the LoG detector option based on brightness and signal-to-noise ratio. Once identified, particles were tracked for their full lifetime using the LAP tracker. This LAP tracker follows molecular displacement as a function of time. Particle trajectories were filtered based on Track Start (removed trajectories that began in first frame), Track End (removed trajectories present in last frame), Duration (removed trajectories ≤ 2 frames and singular extra-long tracks), Track displacement (removed immobilized particles displacement <0.1), and X -Y location (removed particles near the edge of the images). The TrackMate output files were analyzed using PRISM 9 (Graphpad) to calculate characteristic dwell times and diffusion coefficients.

Single exponential curve fit:

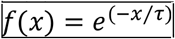

Two exponential curve fit:

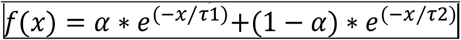

To calculate the diffusion coefficient (μm^2^/sec), we plotted probability density (i.e. frequency divided by bin size of 0.01 μm) versus step size (μm). The step size distribution was fit to the following models:

Single species model:

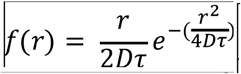

Two species model:

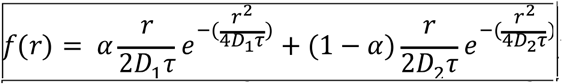

#### Image analysis, curve fitting, and statistics

Image analysis was performed on ImageJ. PRISM 9 (GraphPad) was used for curve fitting. Single molecule dwell time and step size presented in this manuscript represent combined data from 3 technical replicates with 2-3 movies acquired from multiple fields of view for each experimental condition. Dwell time distributions and curve fits were generated with n = 1000-3000 particle trajectories. Step size distribution plots and curve fits represent 10,000-30,000 measured displacements.

